# Fragment-based screening targeting an open form of the SARS-CoV-2 main protease binding pocket

**DOI:** 10.1101/2023.11.15.567102

**Authors:** Chia-Ying Huang, Alexander Metz, Roland Lange, Nadia Artico, Céline Potot, Julien Hazemann, Manon Müller, Marina Dos Santos, Alain Chambovey, Daniel Ritz, Deniz Eris, Solange Meyer, Geoffroy Bourquin, May Sharpe, Aengus Mac Sweeney

## Abstract

To identify starting points for therapeutics targeting SARS-CoV-2, the Paul Scherrer Institute and Idorsia decided to collaboratively perform an X-ray crystallographic fragment screen against its main protease. Fragment-based screening was carried out using crystals with a pronounced open conformation of the substrate binding pocket. Of 631 fragments soaked, a total of 29 hits bound either in the active site (24 hits), a remote binding pocket (2 hits) or at crystal packing interfaces (3 hits). Notably, two fragments with a pose sterically incompatible with a more occluded crystal form were identified. Two isatin-based electrophilic fragments bound covalently to the catalytic cysteine residue. Our structures also revealed a surprisingly strong influence of the crystal form on the binding pose of three published fragments used as positive controls, with implications for fragment screening by crystallography.

**Synopsis:** An X-ray crystallographic screen on SARS-CoV-2 3CL protease resulted in 29 fragment hits, including two isatin-based reversible covalent binders, and revealed a strong influence of the crystal form used for fragment soaking on the bound conformation of three additional reference fragments.

## 1. Introduction

Identifying new chemical leads is a key step in the search to find small-molecule drugs. Fragment-based drug discovery (FBDD) has been developed over the last 20 years and it has been increasingly used in drug discovery, especially by the pharmaceutical industry (Knight *et al*., 2022). The approach uses a range of different methods to screen fragments (low molecular weight ligands obeying the “rule of three”) (Jhoti *et al*., 2013) against a relevant biological target. With the development of automation and higher throughput, X-ray crystallography has become an efficient method to detect weakly binding fragments and directly identify their binding mode to the target protein.

SARS-CoV-2 has killed millions of people and wreaked havoc on the global economy since its fast spread in 2019. Despite the development of vaccines and small molecule drugs it remains a significant global health burden (Msemburi *et al*., 2023). The spread of new variants further increases the medical need for antiviral therapeutics. SARS-CoV-2 encodes several accessory proteins, 4 structural proteins and 16 non-structural proteins (NSP), of which the 3CL protease (main protease, NSP5, 3CL^pro^) is the best studied and essential protease for processing SARS-CoV-2 polyproteins (Jin *et al*., 2020, Zhang *et al*., 2020). The reversible covalent 3CL^pro^ inhibitor PF-07321332 (nirmatrelvir) is used in combination with ritonavir as Paxlovid, an approved treatment for SARS-CoV-2 (Owen *et al*., 2021). Additionally, several inhibitors of 3CL^pro^ and other targets are undergoing clinical trials (Mukae *et al*., 2022, Lei *et al*., 2022). Many of these inhibitors emerged through repurposing, e.g. of inhibitors of proteases (Pang *et al*., 2023). In contrast, fragment screening and FBDD inherently aim at identifying novel starting points.

FBDD studies have identified hundreds of fragments that bind to 3CL^pro^ (Gunther *et al*., 2021, Douangamath *et al*., 2020). The COVID Moonshot project (Boby *et al*., 2023, Chan *et al*., 2021, Zaidman *et al*., 2021) has generated a wealth of small molecules, inhibition data and inhibitor complex crystal structures. To contribute to the challenging global hit finding effort, we used a crystal form of 3CL^pro^ with an active site conformation that is more open and less restricted by crystal packing than that used in most other crystallographic studies of 3CL^pro^ (Gunther *et al*., 2021, Douangamath *et al*., 2020) (**Fig. S1**) to screen 631 fragments using the Fast Fragment and Compound Screening (FFCS) facility at the Swiss Light Source (SLS) (Kaminski *et al*., 2022, Stegmann *et al*., 2023). This open crystal form has been reported previously (Zaidman *et al*., 2021, Sutanto *et al*., 2021) but was obtained only by co-crystallization with potent, covalent inhibitors, and was not previously used for fragment screening. With an overall hit rate of 4.5% (similar to that reported by Covid Moonshot), this crystal based fragment screening campaign identified 29 novel fragments in the active site and remote pockets of 3CL^pro^.

Notably, the fragment screening hits include 3CL^pro^ structures of two fragments based on the reversible, covalent binding motif isatin (indoline-2,3-dione). Isatin is both a known covalent inhibitor of cysteine proteases (Badavath *et al*., 2022, Cheke *et al*., 2022, Jiang & Hansen, 2011, Webber *et al*., 1996, Zhou *et al*., 2006) and an endogenous compound that is present at concentrations of 0.1-10 µM in human tissue (Medvedev *et al*., 2007) and the gut microbiome (Medvedev & Buneeva, 2022). Twenty isatin derivatives have been reported by the COVID Moonshot consortium (Morris *et al*., 2021), with RapidFire assay IC_50_ values as low as 740 nM (SMILES code Cc1nc(CN2C(=O)C(=O)c3cc(Br)ccc32)cs1, Moonshot submission number: LOR-NOR-c954e7ad-1, https://covid.postera.ai/). Structures of seven isatin inhibitors bound to 3CL^pro^ were reported on the Fragalysis site and have just been released in the PDB (Boby et al., 2023) at the time of writing. Isatin-based inhibitors have been modelled in the 3CL^pro^ active site as both covalent (Bao *et al*., 2023) and non-covalent inhibitors (ElNaggar *et al*., 2023, Badavath *et al*., 2022, Liu *et al*., 2020), and the structures and surface plasmon resonance (SPR) results reported in this paper confirm a reversible, covalent binding mode.

Moreover, despite the overlap with results from other fragment screening campaigns, we report several new motifs and interactions. In conclusion, the fragment screening results reported in this study provide further information about potential chemical starting points for inhibitors of this pharmaceutically important target.

## 2. Materials and Methods

### 2.1. Cloning, protein expression and purification of SARS-CoV-2 3CL^pro^

DNA encoding a recombinant fusion protein (supplementary information) composed of N-terminal hexa-histidine tagged SUMO and 3CL^pro^ (NC_045512.2, Nsp5, YP_009742612, Wuhan-Hu-1) was codon optimized for expression in *E. coli* and synthesized (GenScript), based on the published 3CL^pro^ expression and crystal structure (Jin *et al*., 2020). The synthetic DNA was cloned into pET29a (+) using the NdeI and BamHI restriction sites (GenScript) and transformed into BL21(DE3) cells. The protein was expressed overnight (Luria broth medium, 25 µg/ml Kanamycin) at 18°C after inducing with 0.5 mM isopropyl-b-d-thiogalactoside (IPTG) at an OD_600_ of approximately 0.7. Overnight cultures were collected by centrifugation and the recovered cell paste was stored at −70°C. Twelve grams of cell paste was resuspended in 20 mM Tris-HCl at pH 7.8, 150 mM NaCl, 5 mM imidazole and treated with lysozyme (1mg/ml; 30 min) and Benzonase (2500 Units, 10 mM MgCl_2_; 15 min, room temperature). Bacterial cells were lysed by high pressure homogenization (29008 p.s.i. or 200 MPa, Microfluidics MP110P, DIXC H10Z) and centrifuged for 30 minutes at 16000 r.p.m. (Fiberlite F21-8×50y, maximum r.c.f. 30,392 *g*). The hexa-histidine SUMO-3CL^pro^ fusion protein was purified by immobilized metal affinity chromatography (IMAC) with a HisTrap column (5 ml, Cytiva) connected to a FPLC AKTA Purifier 100 system. Histidine tagged fusion protein was eluted with a linear gradient of increasing imidazole concentration (from 0 to 100% elution buffer over 20 column volumes; elution buffer: 20 mM Tris-HCl pH 7.8, 150 mM NaCl, 500 mM imidazole). Eluate fractions containing the target protein were combined and concentrated (Amicon, 10 kDa cutoff). The fusion protein was treated with SUMO protease (Sigma-Aldrich SAE0067, 5 U/ mg target protein) to liberate 3CL^pro^ with authentic N- and C-termini (Ser1 and Gln306, respectively). The mixture of cleavage products was dialyzed overnight at 4°C using a Slide-A-Lyzer cassette (10 kDa cutoff, Thermo Scientific) in 4 l dialysis buffer (20 mM Tris-HCl, 150 mM NaCl). The histidine tagged SUMO protein was separated from non-tagged authentic 3CL^pro^ present in the dialysate by immobilized-metal affinity chromatography (IMAC), collecting 3CL^pro^ in the flow through. 3CL^pro^ was further purified by size exclusion chromatography (HiLoad 26/600 Superdex 200) with storage buffer (20 mM Tris-HCl, 150 mM NaCl, 1 mM TCEP, 1 mM EDTA). The elution volume of 3CL^pro^ indicated a dimer as the oligomeric state. 3CL^pro^ (97% purity by LC-MS analysis) was concentrated (Amicon, 10 kDa cutoff) to a final protein concentration of 26 mg/ml (520 µM) and stored at −70°C.

A 3CL^pro^ variant carrying a C-terminal Avi tag (G_307_SGLNDIFEAQK_318_IEWHE) was produced in the same way to the recombinant wild type protein. To prevent autocleavage of the Avi tag, Gln306 of Mpro has been replaced by a glutamate (Q306E variant). Western blot analysis with a streptavidin-HRP conjugate confirmed biotinylation (K_318_) of 3CL^pro^ Q306E during expression in the *E. coli* host strain mediated by bacterial cell endogenous BirA Ligase. Biotinylated Mpro was used for tethering to SPR sensor chips for small molecule interaction analysis.

### 2.2. FRET-based 3CL^pro^ proteolytic activity assay

The enzymatic activity of the recombinant SARS-CoV-2 main protease 3CL^pro^ was determined by a fluorescence resonance energy transfer (FRET) assay using a custom synthesized peptide substrate with (7-Methoxycoumarin-4-yl)acetyl (MCA) as fluorophore and 2,4-Dinitrophenyl (DNP) as fluorescence quencher: MCA-Ala-Val-Leu-Gln-Ser-Gly-Phe-Arg-Lys(Dnp)-Lsy-NH_2_-trifluoroacetate salt (Bachem AG, Bubendorf CH). This peptide substrate amino acid sequence corresponds to the nsp4/nsp5 (3CL^pro^) cleavage site. A substrate stock solution (10 mM) was prepared in 100% DMSO. 40 µl of a 4 µM substrate solution prepared in H_2_O/TWEEN-20 0.01%) is added to a solution (40 µl) containing 3CL^pro^ to start the enzymatic reaction. The final concentrations of the assay reaction ingredients (80 µl) are 5 nM 3CL^pro^, 2 µM peptide substrate (K_m_ 3.17 µM), 1 mM DTT, 1.2% DMSO, 0.01% TWEEN-20, 25 mM TRIS pH 7.4, 0.5 mM EDTA. 3CL^pro^ was diluted (10 nM) from aliquots stored as stock solution (512 µM, −80°C, storage buffer) in 3CL^pro^ assay buffer (50 mM TRIS pH 7.4, 1 mM EDTA, 2 mM DTT, 0.01 % TWEEN-20). The rate of 3CL^pro^ enzymatic activity (v) was determined by monitoring the increase in fluorescence intensity of reactions at room temperature in black microplates (NUNc 384-well F-bottom) with an Infinite M-100 plate reader (Tecan) using 325 nm and 400 nm as wavelengths for excitation and emission, respectively. Test compounds were dissolved in DMSO and screened first at 25 µM final concentration. 3-fold serial dilutions (125 µM – 6.35 nM) of small molecule test compounds are added to determine inhibitory potency. IC_50_ values were determined by an in-house evaluation tool (IC_50_ Studio with 4-parametric fitting, Hill-equation).

### 2.3. Crystallization of SARS-CoV-2 3CL^pro^

Aliquots of purified 3CL^pro^ at 26 mg/ml in storage buffer were thawed on ice and incubated for 16-18 hours at 20°C with a 10-fold molar excess of the inhibitor GC376 (Fu *et al*., 2020, Ma *et al*., 2020). Vapor diffusion crystallization trials were performed at 20°C using the Morpheus^®^ crystallization screen (MD1-46, Molecular Dimensions) with sitting drops containing 300 nl each of protein and precipitant solution (IntelliPlate 96-2, Art Robbins). A single crystal was grown using 300 mM sodium nitrate, 300 mM disodium hydrogen phosphate, 300 mM ammonium sulphate, 100 mM MES/imidazole pH 6.5, 10% (w/v) PEG550 MME and 20% (w/v) PEG 20K (Morpheus^®^ condition C1) as precipitant. The crystal was then crushed in the sample well and transferred into a seed bead tube (Hampton Research) to obtain a homogeneous suspension of seeds. These seeds were used to crystallize 3CL^pro^ in the absence of the inhibitor GC376, to generate the same crystal form (“Type 1”, space group C2 with cell dimensions matching PDB entry 7c6u). This second round of seeding was considered essential to prevent contamination of the final crystals by the potent inhibitor GC376. Interestingly, although CG376-free Type 1 seeds could be generated, their use in the absence of an inhibitor resulted in a different crystal form (“Type 2”, with space group *P*2_1_2_1_2_1_ and matching the PDB entry 7lcr)(Vuong *et al*., 2021). As Type 1 crystals could not be grown easily without an inhibitor, the Type 2 crystal form was evaluated and selected for subsequent crystallization experiments.

For reproducible, large-scale crystallization using crystal seeds, 3CL^pro^ was diluted to 8 mg/ml in 20 mM Tris pH 7.8, 150 mM NaCl, 1 mM TCEP, 1 mM EDTA and 3 mM DTT. DTT was added because soaking of a small test set of fragments in the absence of a reducing agent revealed oxidation of the active site cysteine during crystallization and soaking. This solution and the seed stock solution produced previously were then used to prepare the large-scale, 631-well crystallization. The crystallization trials were set up by transferring 600 nl protein solution and 50 nl seed stock onto an SwissCI 3-lens crystallization plate (SWISSCI) and adding 550 nl of the original crystallization condition described above using a Mosquito robotic dispenser (SPT Labtech). The plates were sealed with ClearVue sealing sheets (Molecular Dimensions), incubated and imaged at 20°C with a Rock Imager 1000 (Formulatrix). Hexagonal plate-like crystals appeared after 1 day and grew to a maximum size of 150 × 80 × 20 µm^3^ after 3 days. These crystals were used within 7 days for fragment soaking.

### 2.4. Fast fragment and compound screening (FFCS)

The FFCS pipeline established at the Swiss Light Source was used to perform the fragment soaking and screening (Kaminski *et al*., 2022, Stegmann *et al*., 2023). To determine the optimal DMSO concentration for fragment soaking, the 3CL^pro^ crystals were first soaked for 3 hours with 10%, 20% and 30% DMSO using an Echo550 acoustic liquid handling robot (Labcyte) and later harvested and measured with X-ray diffraction. A soaking concentration of 20% DMSO was selected, based on the results of X-ray diffraction which showed no deterioration of the data quality.

For crystal based fragment screening, 631 fragments at 100 mM concentration in DMSO were prepared in either Echo Qualified 384 low dead volume COC microplates (Beckman Coulter) or Echo Qualified 384 well polypropylene microplates (Beckman Coulter), and they were then acoustically dispensed into SwissCI 3-lens crystallization plates (SWISSCI) at a final fragment concentration of 20 mM (20% final concentration of DMSO) using an Echo550 system (Labcyte). The plates were then sealed and incubated at 20°C. The fragment-soaked crystals were harvested after 3 hours, and the Crystal Shifter robot (Oxford Lab Technologies) was used to facilitate and record the harvesting process. The crystals were harvested using MiTeGen cryoloops and snap-cooled in liquid nitrogen without further added cryoprotectant. The loop-mounted samples were placed in Unipucks for X-ray data collection.

### 2.5. Data collection, processing and structural determination

X-ray diffraction experiments were carried out on protein crystallography beamline X06SA-PXI at the Swiss Light Source (SLS), Villigen, Switzerland. Data were collected at 100 K using a cryo-cooled loop in a cryo-stream. Measurements were made using the Smart Digital User (SDU) (Smith *et al*., 2023) developed at the SLS with crystal rotation steps of 0.2° at a speed of 0.01 s/step using an EIGER 16M detector (Dectris) operated in continuous/shutterless data collection mode. The beam transmission, flux and beam size were 40%, ∼1.1 × 10^11^ photons s^−1^ and 60 × 40 µm^2^, respectively. The estimated X-ray dose was 5 MGy for a 360° data set. The data were processed and scaled using AutoProc (Vonrhein *et al*., 2011) and XSCALE (Kabsch, 2010), respectively. Automated molecular replacement was carried out using Dimple (Collaborative Computational Project, 1994) with the 3CL^pro^ structure as template, and PanDDA (Pearce *et al*., 2017) was used for the automated detection and analysis of weakly bound fragments. *Coot* (Emsley *et al*., 2010) was used for model building. *Phenix.refine* (Liebschner *et al*., 2019), BUSTER (Liebschner *et al*., 2019) and REFMAC (Murshudov *et al*., 2011) were used for refinement of the structures. Data collection and refinement statistics are reported in **Table S1**. Figures of molecular structures were generated with *PyMOL* (Schrodinger, 2015). Ligand restraints were generated using Pyrogen (Collaborative Computational Project, 1994) or Grade2 (Smart, 2021), using the covalently bound chemical structure for cpds 27-29.

Similar measurements were carried out at a wavelength of 2.066 Å (6 keV) to locate DMSO molecules using the anomalous signal of sulfur. The beam transmission, flux and beam size were 30%, ∼6.6 × 10^10^ photons s^−1^ and 60 × 40 µm^2^, respectively. The estimated dose was 375 K Gy for a 360° data set.

All diffraction data and refined models have been deposited in the Protein Data Bank (PDB) with PDB Entry IDs (7gre, 7grf, 7grg, 7grh, 7gri, 7grj, 7grk, 7grl, 7grm, 7grn, 7gro, 7grp, 7grq, 7grr, 7grs, 7grt, 7gru, 7grv, 7grw, 7grx, 7gry, 7grz, 7gr0, 7gr1, 7gr2, 7gr3, 7gr4, 7gr5 and 7gr6) as listed in **Table S1**. The SMILES codes of the 29 fragments are reported in **Table S2.**

### 2.6. Surface plasmon resonance (SPR)

SPR experiments were performed using a Biacore T200 equipped with a Series S Sensor Chip SA. Biotinylated MproQ306E was immobilized to the streptavidin covalently attached to a carboxymethyl dextran matrix. The initial conditioning of the surfaces on flow cell 1 and 2 was performed by three 1-minute pulses of 1 M NaCl, 50 mM NaOH solution. The ligand at a concentration of 0.27 mg/ml in immobilization buffer (10 mM HEPES, 150 mM NaCl, 1mM TCEP, 0.05% polyoxyethylene (20) sorbitan monolaurate (P20), pH 7.4) was immobilized at a density of 3000 RU on flow cell 2 at a flow rate of 5 μl/min and flow cell 1 was left blank to serve as a reference surface. Surfaces were stabilized with 3 hours injection at a flow rate of 40 µL/min of running buffer (10 mM HEPES, 150 mM NaCl, 1mM TCEP, 0.05% P20, 5% DMSO, pH 7.4).

To collect binding data, sample in 10 mM HEPES, 150 mM NaCl, 0.05% P20, 5% DMSO, pH 7.4, was injected over the two flow cells at 625 µM at a flow rate of 40 μl/min and at a temperature of 25°C. The complex was allowed to associate and dissociate for 50 and 100 s, respectively for each sample.

A DMSO correction curve was performed before/after every 104 cycles. Data were collected at a rate of 10 Hz analyzed within Biacore T200 Evaluation software. 6-Chloro-chroman-4-carboxylic acid isoquinolin-4-ylamide was used at 1.25 µM as a positive control. The structure of this inhibitor bound to 3CL^pro^ has previously been solved as Fragalysis entry P0012_0A:ALP-POS-CE760D3F-2 (https://fragalysis.diamond.ac.uk/).

## 3. Results and Discussion

Automated analysis of the fragment soaked structures using PanDDA (Pearce *et al*., 2017) was followed by visual inspection of potential binding events and full refinement of approximately 60 selected structures. This resulted in a final set of 29 novel fragment complex structures, including three with borderline electron density for the fragment and three covalently bound compounds. An overview of the fragment binding pockets and electron density maps are shown in **Figs. 1 and 2, respectively**, with the binding site described using Schechter and Berger notation (Schechter & Berger, 1967) (**Fig. S2**). Twenty-four of the fragments were bound in the substrate binding pocket, while two fragments were observed in a remote pocket close to the C-terminus (**Fig. 1**). The remaining three fragments occupied two different remote binding sites partly formed by crystal contacts. Due to their location at crystal packing interfaces, these compounds (cpds 24-26) were considered as potential artefacts.

**Figure 1.**
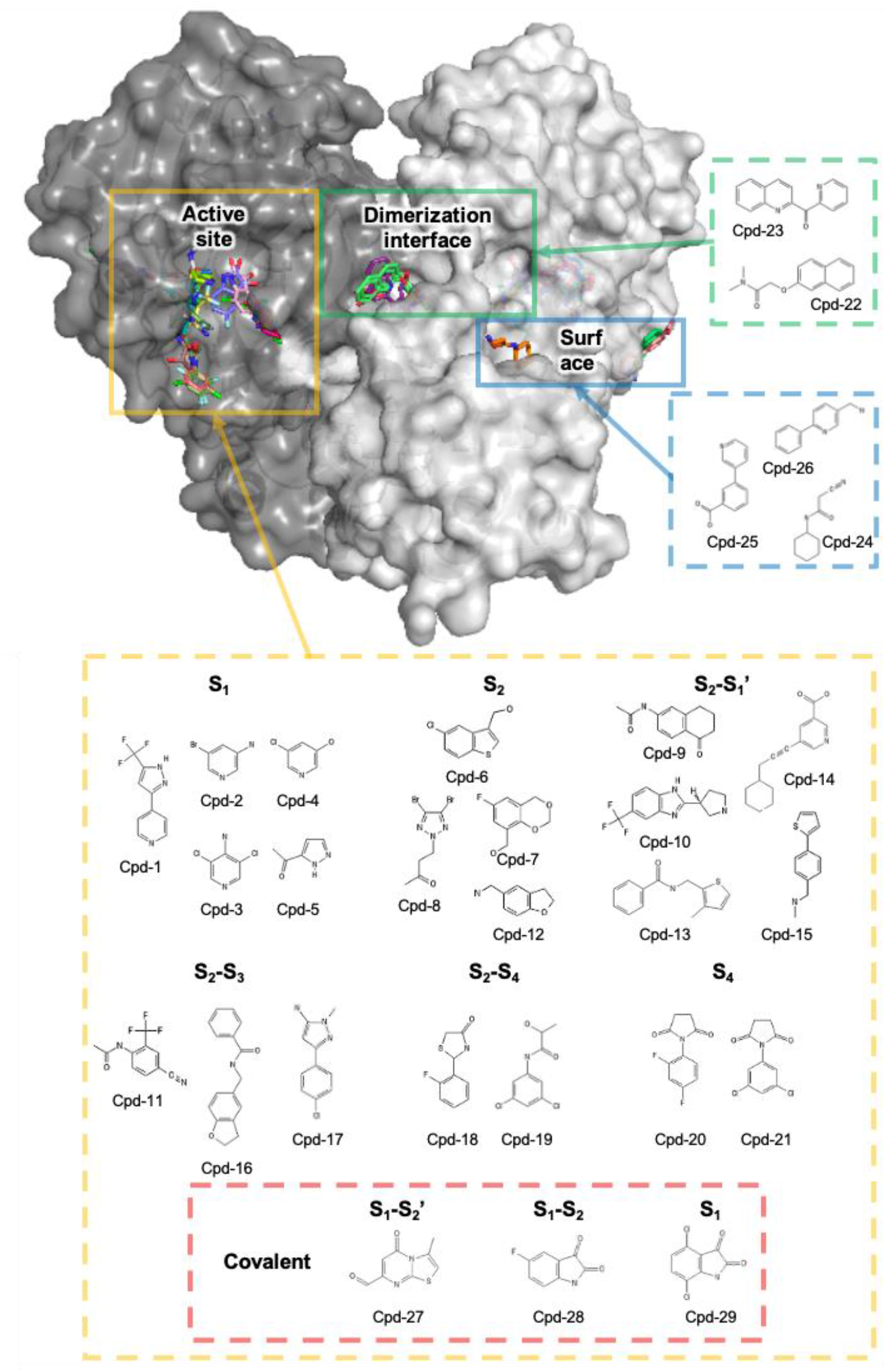
Overall structure of 3CL^pro^ showing the 29 fragment hits. For clarity, only one substrate binding pocket of the homodimer is shown. The protein is shown in surface representation and fragments are shown in stick representation.

**Figure 2.**
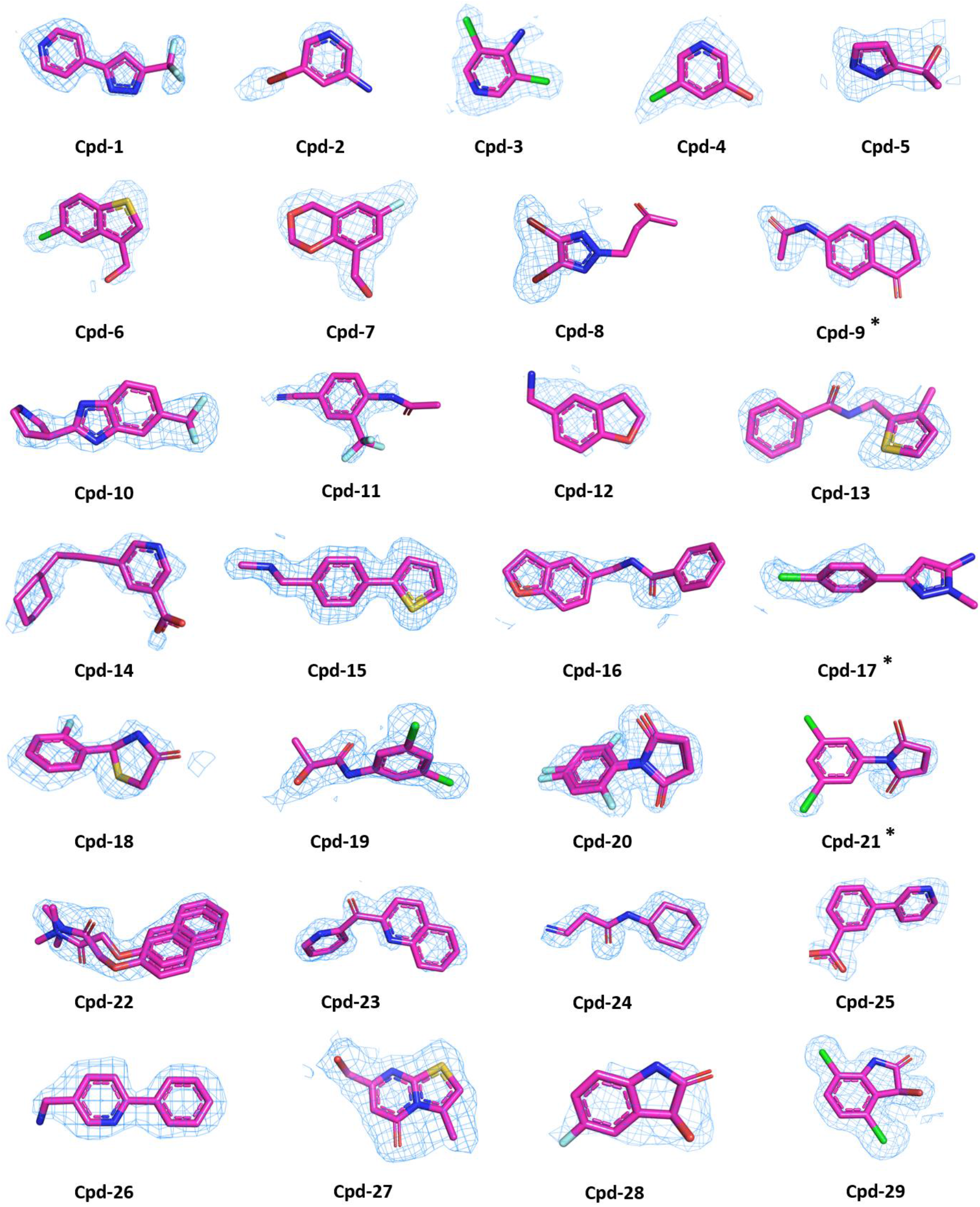
Final 2Fo-Fc electron density for all fragments. A 1.0 sigma contour is shown in blue except for cpd-9, cpd-17 and cpd-21 (marked with *), which are shown at 0.7 sigma contour. Density within 2 Å of the ligand molecule is shown. Stick models show C (magenta), N (blue), O (orange red), Br (dark red), S (yellow), Cl (green) and F (light blue) atoms.

### 3.1. Fragment binding sites and interactions

Representative active site binders were selected according to their binding mode. The binding of one representative fragment is described for each of the additional binding sites.

#### 3.1.1. Active site subpockets

The published structure (PDB entry 7mgr) of the NSP 8/9 substrate peptide in complex with the inactive Cys145Ala variant of 3CL^pro^ is used to denote the S_1_-S_4_ pockets. The peptide residue Gln5 is bound in S_1_, with Leu4 in S_2_, Lys3 in S_3_ and Val2 in S_4_. Where possible, active site figures match the standard protease orientation as shown in **Fig. S2**. In this orientation, the N-terminal residues of a peptide substrate are shown on the left and the C-terminal residues on the right.

#### 3.1.2. Open and flexible active site

The crystal form used for these soaking experiments has a more open conformation than, for example, the widely used *C*2 crystal form (PDB entry 5r83, Douangamath *et al*., 2020) (**Fig. S1**). The C_α_ atoms of chain A residues 43-52 are shifted by 1.0-1.8 Å from the corresponding residues in PDB entry 5r83, with the Met49 and Ser46 side chains shifted by 3.0 Å and 2.8 Å, respectively. 3CL^pro^ is present as a homodimer in the asymmetric unit in our structures, and as a monomer in the asymmetric unit in the *C*2 form, with the crystallographic two-fold axis relating both monomers of the homodimer. As the flexible 3CL^pro^ active site samples many conformations in solution, we believe that the use of different, complementary crystal forms for fragment screening maximizes the diversity of chemical starting points. Furthermore, the flexibility of the open binding pocket is less restricted by crystal contacts.

### 3.2. Non-covalent active site binders

#### 3.2.1. S_1_ pocket

Cpd-2 is bound in the S_1_ pocket (**Fig. 3a)**. The high scattering power of the Br atom allowed the identification of this very small fragment, which is bound both in molecule A and B. In structures with different fragments, we found alternative occupation of the S_1_ pocket by DMSO and the overlapping fragment. In this case, long wavelength X-ray data collection at 6 keV was also carried out to check the DMSO occupancy at this location. The absence of a sulphur anomalous peak provided a further strong indication of the cpd-2 bound 3CL^pro^ complex.

**Figure 3.**
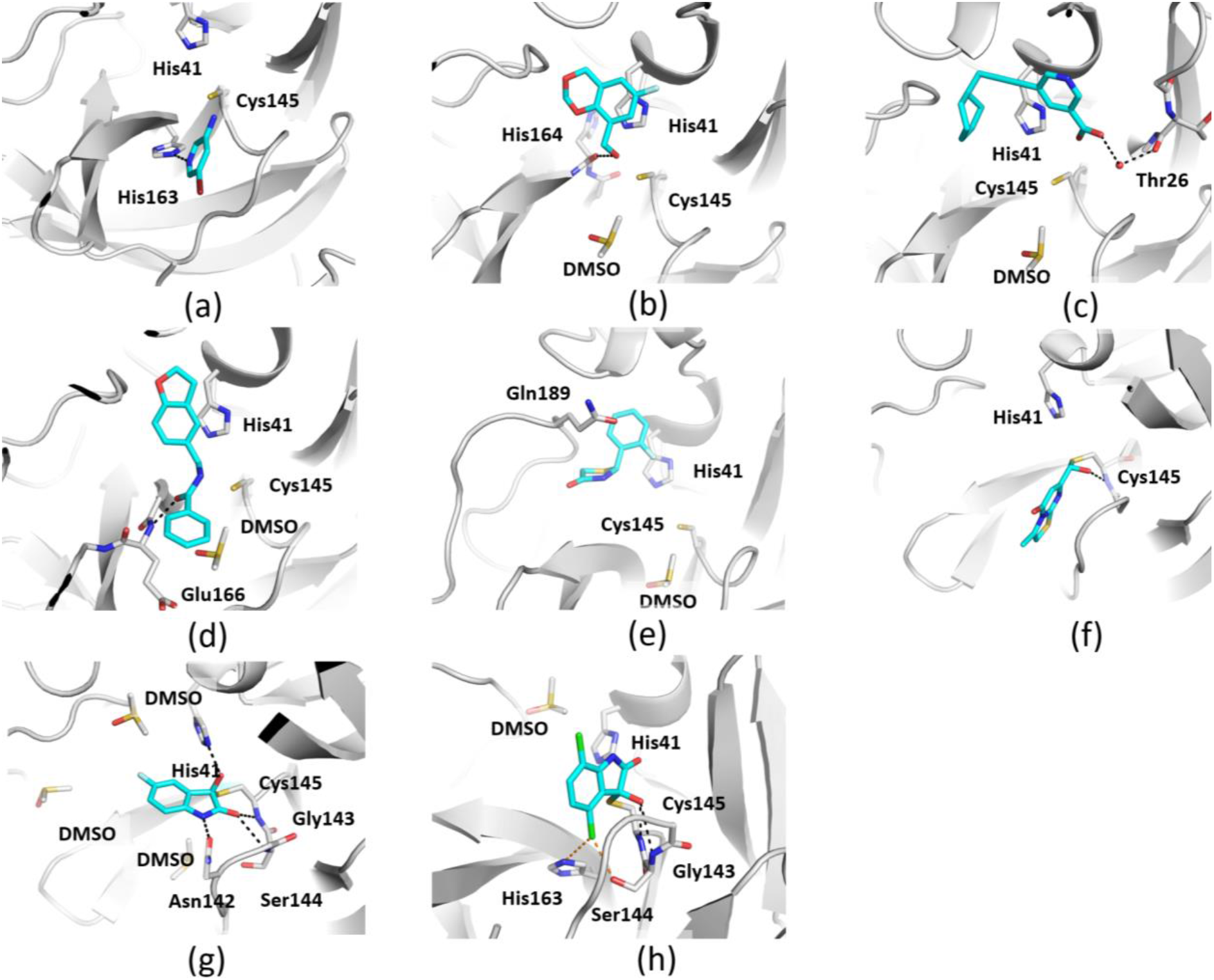
The active site bound fragments (a) cpd-2, (b) cpd-7, (c) cpd-14, (d) cpd-16, (e) cpd-18, (f) cpd-27, (g) cpd-28 and (h) cpd-29. Hydrogen bonds are shown as black dashes. Halogen bonds are shown as orange dashes in (f). The color code for the stick representation of the fragment is the same as described in Fig. 2 with the carbon in cyan.

Cpd-2 forms a hydrogen bond with His163 and an interaction with Cys145(S) and adjacent water. A substructure search of the PDB using cpd-2 did not identify any ligands. Searching with the Br atom excluded identified the known fragment hit (PDB entry 5re4), with a similar binding pose and hydrogen bond with His163 but with the plane of the ring rotated by approximately 30 degrees. For the remaining compounds, substructure searches and similarity searches using the PDB query tool revealed no similar 3CL^pro^ ligands. Other fragments bound to the S_1_ pocket are cpds-1, 3, 4 and 5. Moreover, cpd-3 and 5 additionally interacts with Asn142(O).

#### 3.2.2. S_2_ pocket

Cpd-7 is bound with the dioxane buried in the S_2_ pocket (**Fig. 3b**). The fragment forms a single hydrogen bond with His164(O), and a face-to-face aryl interaction with His41. The hydroxy group acts as a hydrogen bond donor, interacting with the His164 backbone carbonyl, but also acts as a hydrogen bond acceptor in water mediated hydrogen bonding. Other fragments bound to the S_2_ pocket are cpd-6, -8 and -12.

#### 3.2.3. S_1_′ pocket

Cpd-14 occupies the S_2_ and S_1_′ pockets, and extends to the entrance of the S_2_′ pocket (**Fig. 3c**). The cyclohexyl fits in the hydrophobic S_2_ pocket. The carboxylic acid forms water mediated hydrogen bond interactions with Thr26 and a putative sulfenic acid state of Cys145. The fragment is bound in molecule A only. Notably, the 3-carboxy-pyridine of this fragment is directed into the S_1_′-region of the substrate binding cleft, which is rarely occupied by other fragments. Such a bridging of the catalytic center by a non-peptidic motif may be interesting for the design of inhibitors simultaneously addressing the prime and non-prime portions of the binding cleft. Positive difference electron density at Cys145 of molecule A (only) indicated a modification of the cysteine sulphur, possibly an oxidation to form the normally unstable species peroxysulfenic acid as described previously for 3CL^pro^ (Kneller et al., 2020).

#### 3.2.4. S_2_ and S_3_ pockets

Cpd-16 extends from S_2_ to S_3_ (**Fig. 3d**), forming a hydrogen bond with Glu166(N) and a face-to-face aryl interaction with His41. The central amide carbonyl forms a hydrogen bond with the backbone NH of Glu166. The phenyl ring forms no binding interactions with the protein and is more mobile than the rest of the fragment. The fragment is bound in molecule A only. Some positive Fo-Fc difference electron density adjacent to Cys145(S) was not modelled and may be due to partial oxidation.

#### 3.2.5. S_2_ to S_4_ pockets

Cpd-18 extends from S_2_ towards S_4_ (**Fig. 3e).** The 2-chlorophenyl forms a face-to-face aryl interaction with His41. Unlike cpd-16, which occupies the S_2_ and S_3_ pockets, the thiazolidinone carbonyl of cpd-18 extends directly towards S_4_, displacing the side chain of Gln189. Cpd-6 also occupies the S_4_ pocket and displaces the Gln189 side chain.

#### 3.2.6. S_4_ pocket

Cpds 20 and 21 bind congruently, with their respective N-linked succinimide portion in the S_4_ pocket (**Figs. 4a and b**). This motif has not been observed previously and forms hydrogen bond interactions with its carbonyl groups to Gln192(NH) and the side chain NH of Gln189. In contrast, the fragments’ substituted phenyl portions form no prominent interactions aside from non-polar contacts to Pro168 and Ala191. This fact may indicate this site as a strong interaction hotspot for the succinimide motif. Furthermore, these fragments bind without utilizing the more affinity-providing subpockets S_1_-S_3_ closer to the active site, yet may possess viable exit vectors towards this region through substitution at the imide N or the carbonyl-adjacent carbon position.

**Figure 4.**
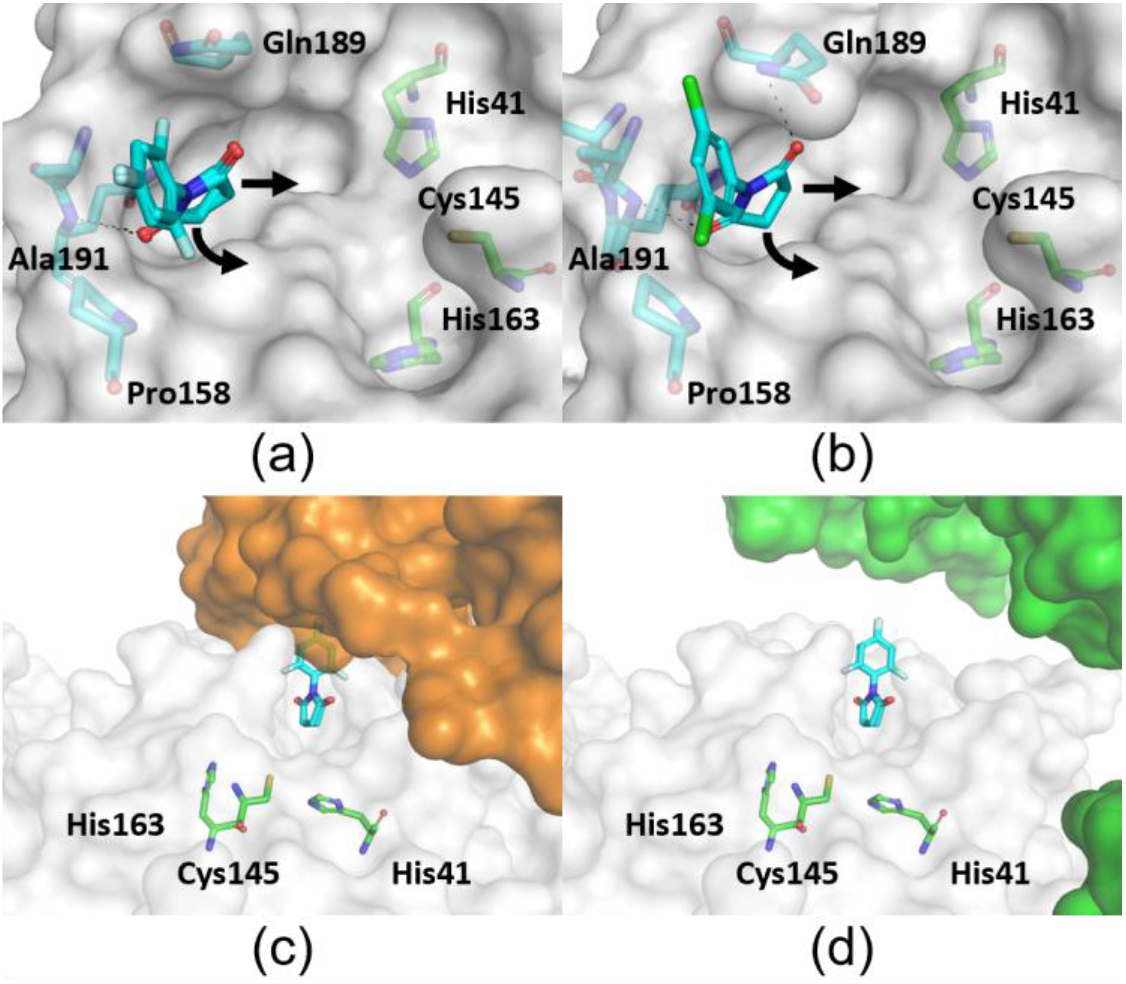
Imide fragments binding and blocked binding cleft in closed crystal form. (a) Cpd-20 and (b) cpd-21 bind congruently, with their respective N-linked succinimide portion in the S_4_ pocket. Potential exit vectors towards the catalytic center indicated by arrows. (c) These fragments would clash with a crystal mate (orange surface; PDB entry 5rgr) in the closed crystal form; structures aligned with respect to the protein monomer binding cpd-20. (d) The more open crystal form used in this study allows unobstructed access to the complete binding cleft (closest crystal mate as green surface).

The more important observation, however, is that this motif could not have been identified using the closed crystal form used in most other crystallographic screens against 3CL^pro^. In this closed crystal form the phenyl portion of both fragments would clash with a crystallographic symmetry mate that partially blocks the N-terminal section of the peptide binding cleft. (**Fig. 4c**). Thus, it may not be surprising that only very few fragments were reported bound to the S_4_ pocket, which then mostly form interactions with the crystal partner, which puts into question the physiological relevance of the observed position. In contrast, the nearest crystal mate of the crystal form used in this study is further away allowing unobstructed access to the complete binding cleft (**Fig. 4d**). Arguably, this crystal mate may also prevent other fragments from accessing the binding cleft even if their pose does not directly clash with that crystal mate.

### 3.3. Covalent active site binders

#### 3.3.1. Isatin-based fragments

Cpds 28 and 29 bind covalently to Cys145 via their reactive isatin groups (**Fig. 3g and h)**. We observed covalent binding of the isatin-based cpds 28 and 29 to the catalytic Cys145 residue in both active sites of the homodimer. Presumably due to the high concentration (20 mM) of isatin used in the soaking experiment, covalent binding of both compounds to Cys44 was also observed in 3CL^pro^ molecule B of the asymmetric unit. Seven structures of isatin-containing fragments bound to 3CL^pro^ were already solved and shared on https://fragalysis.diamond.ac.uk/, and they have just been released in the PDB (Boby et al., 2023). Interestingly, a structure of an unrelated COVID-19 protein, the NSP3 macrodomain component of the replication complex, with a non-covalently bound isatin molecule has also been published (PDB entry 5rtf, with PanDDA event maps deposited at https://fragalysis.diamond.ac.uk/) (Schuller *et al*., 2021).

The reversible binding mode of both inhibitors was confirmed by a FRET-based biochemical assay (**Table 1**) and by SPR (**Fig. 5**). Similar half maximal inhibitory concentration (IC_50_) values were observed after 30 and 60 minutes of pre-incubation of the enzyme and inhibitor, further supporting a reversible binding mode (as the IC_50_ of irreversible inhibitors decreases with increasing preincubation time).

**Figure 5.**
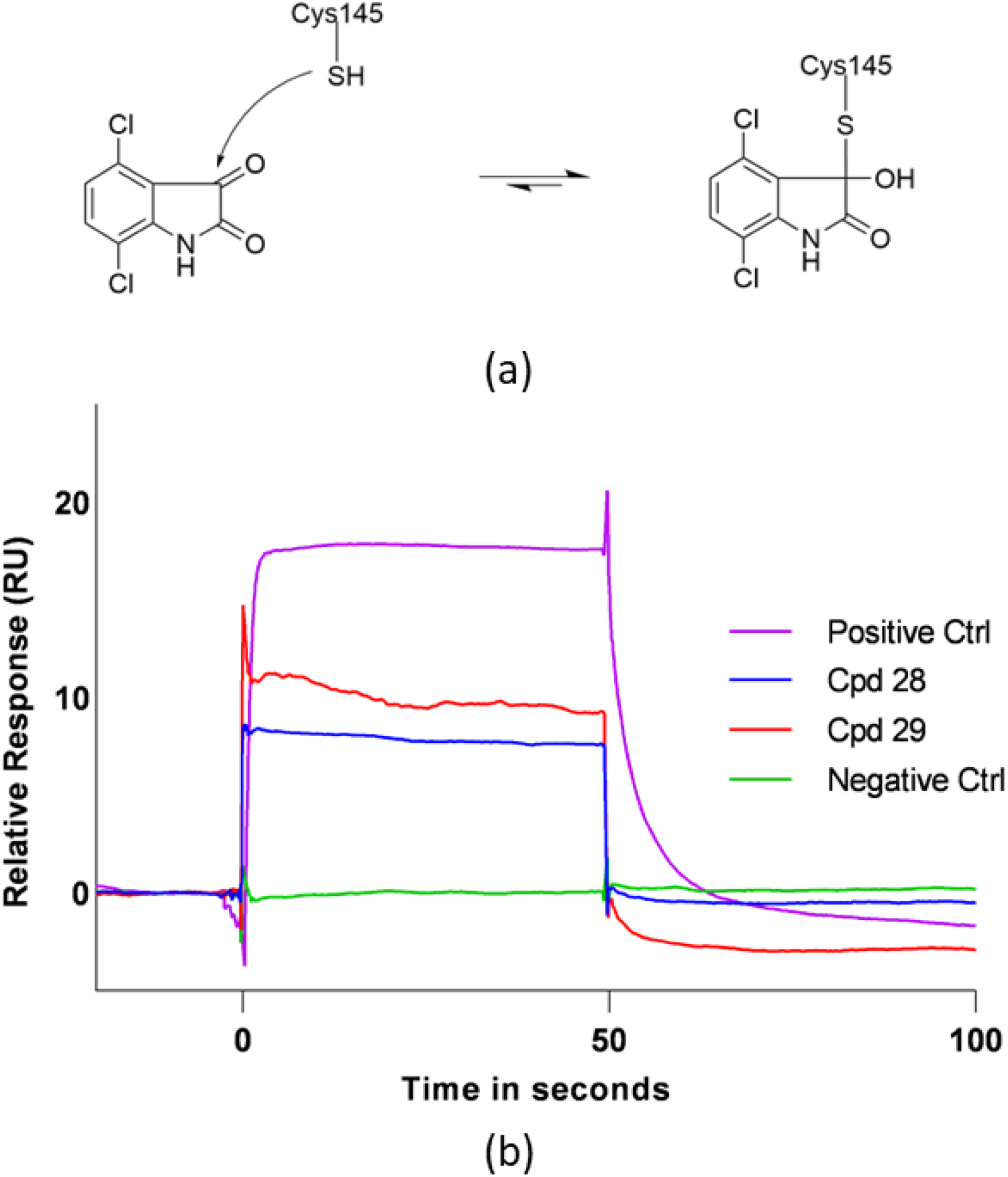
Reversible, covalent inhibition mechanism of 3CL^pro^ by isatin-based inhibitors. (a) The reversible covalent bond formation between an isatin and Cys145. (b) A surface plasmon resonance sensorgram showing rapidly reversible binding of cpds 28 and 29. 6-Chloro-chroman-4-carboxylic acid isoquinolin-4-ylamide was used at 1.25 µM as a positive control, and running buffer containing no 3CL^pro^ fragment was used as a negative control.

**Table 1.**
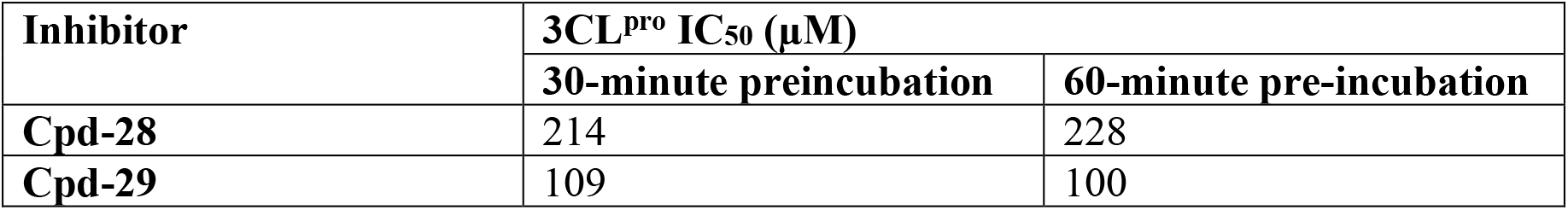
IC_50_ values of compounds 28 and 29 after 30- and 60-minute pre-incubation.

The isatin derivatives cpd-28 and cpd-29 adopt different binding poses. Cpd-29 binds in the S_1_ pocket and forms hydrogen bond interactions with the backbone nitrogen of Gly143 and Cys145, while cpd-28 extends towards S_2_ and forms hydrogen bonds with the backbone of Gly143 and the side chain of Asn142. The bulky chlorine atom on carbon atom 4 of cpd-29 (atoms CL1 and C6, respectively) prevents it from adopting the same binding pose as cpd-28 due to a steric clash with the backbone carbonyl of His164. One of the two chlorine atoms of cpd-29 forms halogen bonds with Ser144 and His163. Covalent binding of isatin to Cys145 creates an R-configured stereo center for both bound inhibitors (**Fig. 5**).

The hydroxy group of the covalently bound cpd-28 forms a hydrogen bond with the His41 side chain. The isatin amino group makes a hydrogen bond interaction with the side chain carbonyl group of Asn142 (**Fig. 3g**).

Structures of isatin covalently bound to monoamine oxidase (PDB entry 1oja) (Binda *et al*., 2003), the cysteine proteases caspase-3 (PDB entry 1gfw) (Lee *et al*., 2000) and rhinovirus 3C protease (Webber *et al*., 1996) and several other proteins have also been published. Isatin derivatives have been published both as covalent and noncovalent inhibitors of SARS (SARS-1) 3CL^pro^ (Zhou *et al*., 2006). Surprisingly, isatin-based inhibitors have been modelled in non-covalent binding poses in the SARS-CoV-2 3CL^pro^ active site and used as the basis for both *in silico* selection of compounds for screening (Badavath *et al*., 2022, Liu *et al*., 2020) and modelling-based inhibitor optimization (ElNaggar *et al*., 2023). The resulting isatin-based inhibitors have been published as potent, non-covalent inhibitors (Jiang *et al*., 2023). The recently deposited structures provided by COVID Moonshot, together with the isatin complex structures and SPR results described here, confirm the reversible covalent binding mode of isatin inhibitors to 3CL^pro^ (**Table 1**). We hope that these structures, as the first SARS-CoV-2 3CL^pro^ structures deposited in the PDB, will provide clear structural evidence of covalent binding and serve as templates for future modelling of covalently bound isatins.

In addition to binding at the active site, cpds 28 and 29 also bound to Cys44 of molecule B. Unlike Cys145, which is activated by its environment, Cys44 does not take part in the catalytic mechanism, and cpd-28 appears to exist as a mixture of non-covalently bound and covalently bound forms in the Cys44 pocket. Non-selective binding of drugs is clearly undesirable and designing a more selective isatin-based 3CL^pro^ inhibitor, for example by tuning the isatin electrophilicity, could be a difficult challenge. Cys44 is close to the substrate binding site, and binding of the isatin inhibitors causes a rearrangement of residues Cys44-Arg60 and Val186-Gly195 and an increase in the size of the S_2_ and S_4_ pockets. In 3CL^pro^ molecule B, this rearrangement would result in steric clashes with the neighboring 3CL^pro^ molecule, and we speculate that this steric hindrance prevents the binding of isatin to Cys44 of 3CL^pro^ molecule A in this crystal form.

While isatins and other covalent inhibitors require careful optimization to selectively inhibit the target of interest, inhibitors that bind covalently to the catalytic Cys145 of 3CL^pro^ may be less prone to resistance development. As this is a concern for small molecule antiviral drugs (DeGrace *et al*., 2022), substrate binding pocket analysis was used to highlight the evolutionarily vulnerable regions of 3CL^pro^ most likely to tolerate mutations leading to resistance development (Shaqra *et al*., 2022). As the catalytic residues of the protease are arguably the most resistant to mutation, we believe that targeting binding through strong interactions with His41 and/or Cys145 of 3CL^pro^ is a promising strategy for the structure guided optimization of robust inhibitors.

#### 3.3.2. Aldehyde-based fragment

Cpd-27 binds in the S_1_ pocket and partially in the S_2_’ pocket while forming a reversible covalent bond with the catalytic Cys145 via its aldehyde group (**Fig. 3f)**. The resulting hemiacetal hydroxy group accepts a hydrogen bond from Cys145(NH) and acts as a hydrogen bond donor forming a water-mediated interaction with Thr26 (not shown in **Fig. 3f**). Aldehydes are known covalent inhibitors of cysteine proteases and peptidomimetic aldehyde inhibitors were among the first described for 3CL^pro^ (Dai *et al*., 2020).

### 3.4. Non-active site binders

3.4.1. Cpds 22 and 23 bind in a pocket that is distinct from the binding site of pelitinib (**Fig. 6a)**, known as the C-terminal dimerization domain (Gunther *et al*., 2021). This cryptic binding pocket is instead formed by repositioning of the residues from Ser301 to the C-terminus. Other ligands observed in this pocket include the fragments 1-methyl-N-{[(2S)-oxolan-2-yl]methyl}-1H-pyrazole-3-carboxamide (PDB entry 5rfa), 1-(4-fluoro-2-methylphenyl)methanesulfonamide (PDB entry 5rgq) (Douangamath *et al*., 2020) as well as PEG (PDB entry 8drz) and DMSO (many structures, including PDB entry 7qt6).

**Figure 6.**
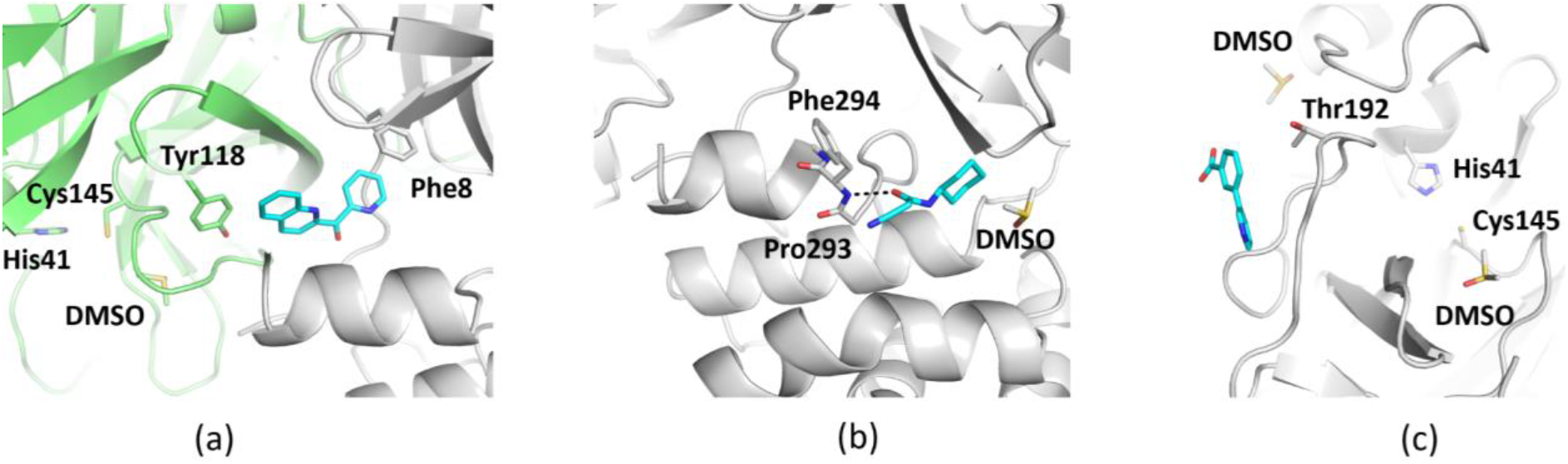
Non-active site binding fragments (a) cpd-23 (b), cpd-24 and (c) cpd-25.

3.4.2. Cpd-24 binds in the same pocket as the compound AT7519 in PDB entry 7aga (**Fig. 6b)**, which is referred to as the second allosteric pocket (Gunther *et al*., 2021). Cpds 25 and 26 bind at the surface of 3CL^pro^ in a pocket that is mainly formed by crystal contacts (**Fig. 6c)**. Other ligands observed at this location include the fragment 4-amino-N-(pyridin-2-yl)benzenesulfonamide (PDB entry 5rf8) (Douangamath *et al*., 2020) and the buffer components PEG (PDB entry 7kvl) and ethylene glycol (PDB entry 7nf5).

### 3.5. Comparison of five reference compounds in two different crystal forms

The fragments from PDB entries 5rh0, 5rh1, 5rh2, 5r83 and 5rgu were used as positive controls during optimization of the crystal soaking conditions. To check for any influence of the more open active site conformation used in this study, the fragment complex structures were compared with the published PDB entries in a different space group (Douangamath *et al*., 2020). Notably, three of the five fragments exhibited significant differences in their binding poses (**Fig. 7)**. While the pyridine moiety of each fragment was fixed via a hydrogen bond to His163, the aromatic ring at the opposite end of each fragment molecule was shifted and rotated by up to 80 ° (**Fig. 7**).

**Figure 7.**
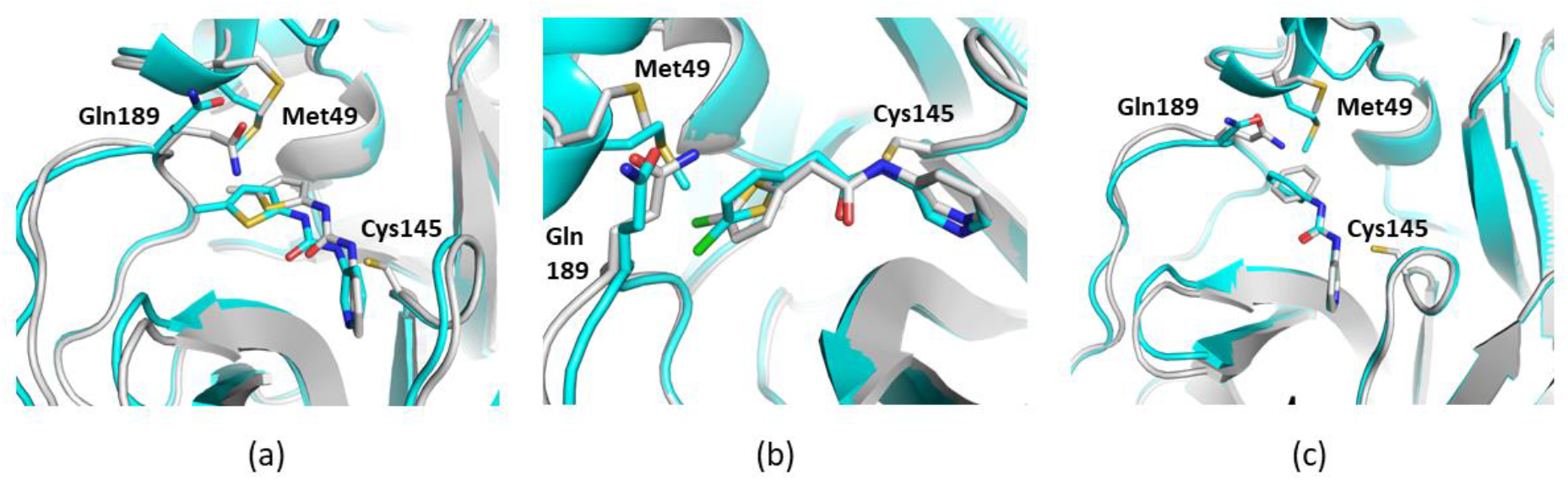
Binding of three fragments in more closed 3CL^pro^ published structures (cyan) compared with this study with a more open 3CL^pro^ conformation (grey). Comparison of the same small molecule in the two different structures shows (a) a translation or (b and c) a rotation of the upper-left ring of fragments corresponding to PDB entries 5rh0, 5rh1 and 5r83, respectively. The position of Met49 shows the more open binding site conformation used in this study.

## 4. Conclusion

The significant differences between the different structures of the same inhibitor show both the advantages and potential risks of soaking fragments into crystals for screening or for structural analysis of validated hits. For flexible binding pockets, restriction of the conformational freedom in the crystal environment may reduce the entropic penalty of binding for a subset of fragments and increase their binding affinity to the crystallized protein. While these hits may bind with a greatly reduced affinity to the free (uncrystallized) target protein, they often provide the first starting points for modeling and chemical exploration. Equally, the reduced flexibility of the binding pocket may prevent it from adopting the conformation required for the binding of other genuine fragment hits. This may partly explain the low correlation between the fragment hits observed when screening by crystallography or other methods (Schiebel *et al*., 2016). Finally, although crystal packing related artefacts can be minimized by co-crystallization, these results highlight the importance of exploring the binding site flexibility and ligand mobility during structure guided drug discovery. Room temperature data collection has been used to probe different active site conformations and avoid cryogenic cooling artefacts (Huang *et al*., 2022).

In summary, we report a crystallographic fragment screen against 3CL^pro^ using a crystal form that is less obstructed by crystal packing and has a more open substrate binding cleft conformation than previously used crystal forms. We identified a number of new or varied motifs and binding interactions with the potential to be instrumental in structure-guided drug discovery. In particular, we identified fragments that could not have been found with more closed crystal forms due to steric overlap, but also revealed varied binding poses of known crystallographic binders in the flexible binding cleft. These observations demonstrate the implications of the chosen crystal form for fragment screening and subsequent structure-guided design. The subsequent use of co-crystallization, different crystal forms and/or room temperature data collection, together with an awareness of the potential influence of the crystal form, may maximize the value of fragment structures in drug discovery projects.

