## Supplementary Information for "Fragment-based screening targeting an open form of the SARS-CoV-2 main protease binding pocket"

Gene construction details

Figures S1 and S2

Table S1 and S2

**Cloning and expression**

SARS-CoV-2 3CL$^{pro}$ expression as a Hexa-histidine-SUMO-fusion protein
amino acid sequence source code :

**MGSSHHHHHHGSGLVPRGSASMSDSEVDQEAKPEVKPEVKPETHINLKVSDGSS
EIFFKIKKTTPLRRLMEAFAKRQGKEMDSLRFLYDGIRIQADQTPEDLDMEDND
IIEAHREQIGGSGFRKMAFPSGKVEGCMVQVTCGTTTLNGLWLDDVVYCPRHV
ICTSEDMLNPNYEDLLIRKSNHNFLVQAGNVQLRVIGHSMQNCVLKLKVDTANP
KTPKYKFVRIQPGQTFSVLACYNGSPSGVYQCAMRPNFTIKGSFLNGSCGSVGF
NIDYDCVSFCYMHHMELPTGVHAGTDLEGNFYGPFVDRQTAQAAGTDTTITVN
VLAWLYAAVINGDRWFLNRFTTTLNDFNLVAMKYNYEPLTQDHVDILGPLSAQ
TGIAVLDMCASLKELLQNGMNGRTILGSALLEDEFTPFDVVRQCSGVTFQ**

Black : Hexa-His tag and linker ; Blue: SUMO ; Red: 3CL$^{pro}$

**Hexahistidine SUMO-3CL$^{pro}$**

425 aas; Mol Wt 47194.9, Isoelectric Pt (pI) 5.80

Amino acid sequence "Backtranseq"; *E. coli* codon usage high
https://www.ebi.ac.uk/Tools/st/emboss_backtranseq/

**DNA synthesized (Genscript) and cloned into pET29a+**
**CAT**ATGGGTTCTTCTCACCACCACCACCACCACGGTTCTGGTCTGGTTCCGCGTG
GTTCTGCGTCTATGTCTGACTCTGAAGTTGACCAGGAAGCGAAACCGGAAGTTAA
ACCGGAAGTTAAACCGGAAACCCACATCAACCTGAAAGTTTCTGACGGTTCTTCT
GAAATCTTCTTCAAAATCAAAAAAACCACCCCGCTGCGTCGTCTGATGGAAGCGT
TCGCGAAACGTCAGGGTAAAGAAATGGACTCTCTGCGTTTCCTGTACGACGGTAT
CCGTATCCAGGCGGACCAGACCCCGGAAGACCTGGACATGGAAGACAACGACAT
CATCGAAGCGCACCGTGAACAGATCGGTGGTTCTGGTTTCCGTAAAATGGCGTTC
CCGTCTGGTAAAGTTGAAGGTTGCATGGTTCAGGTTACCTGCGGTACCACCACCC
TGAACGGTCTGTGGCTGGACGACGTTGTTTACTGCCCGCGTCACGTTATCTGCAC
CTCTGAAGACATGCTGAACCCGAACTACGAAGACCTGCTGATCCGTAAATCTAAC

CACAACTTCCTGGTTCAGGCGGGTAACGTTCAGCTGCGTGTTATCGGTCACTCTA
TGCAGAACTGCGTTCTGAAACTGAAAGTTGACACCGCGAACCCGAAAACCCCGA
AATACAAATTCGTTCGTATCCAGCCGGGTCAGACCTTCTCTGTTCTGGCGTGCTA
CAACGGTTCTCCGTCTGGTGTTTACCAGTGCGCGATGCGTCCGAACTTCACCATC
AAAGGTTCTTTCCTGAACGGTTCTTGCGGTTCTGTTGGTTTCAACATCGACTACGA
CTGCGTTTCTTTCTGCTACATGCACCACATGGAACTGCCGACCGGTGTTCACGCG
GGTACCGACCTGGAAGGTAACTTCTACGGTCCGTTCGTTGACCGTCAGACCGCGC
AGGCGGCGGGTACCGACACCACCATCACCGTTAACGTTCTGGCGTGGCTGTACGC
GGCGGTTATCAACGGTGACCGTTGGTTCCTGAACCGTTTCACCACCACCCTGAAC
GACTTCAACCTGGTTGCGATGAAATACAACTACGAACCGCTGACCCAGGACCAC
GTTGACATCCTGGGTCCGCTGTCTGCGCAGACCGGTATCGCGGTTCTGGACATGT
GCGCGTCTCTGAAAGAACTGCTGCAGAACGGTATGAACGGTCGTACCATCCTGG
GTTCTGCGCTGCTGGAAGACGAATTCACCCCGTTCGACGTTGTTCGTCAGTGCTC
TGGTGTTACCTTCCAGTAATAG**GGATCC**

**Storage buffer (X-ray)**
20 mM Tris-HCl pH 7.8, 150 mM NaCl, 1 mM TCEP, 1 mM EDTA

**Storage buffer (Rapidfire MS)**
20 mM Tris-HCl pH 7.8, 150 mM NaCl, 1 mM TCEP, 1 mM EDTA, 5% glycerol

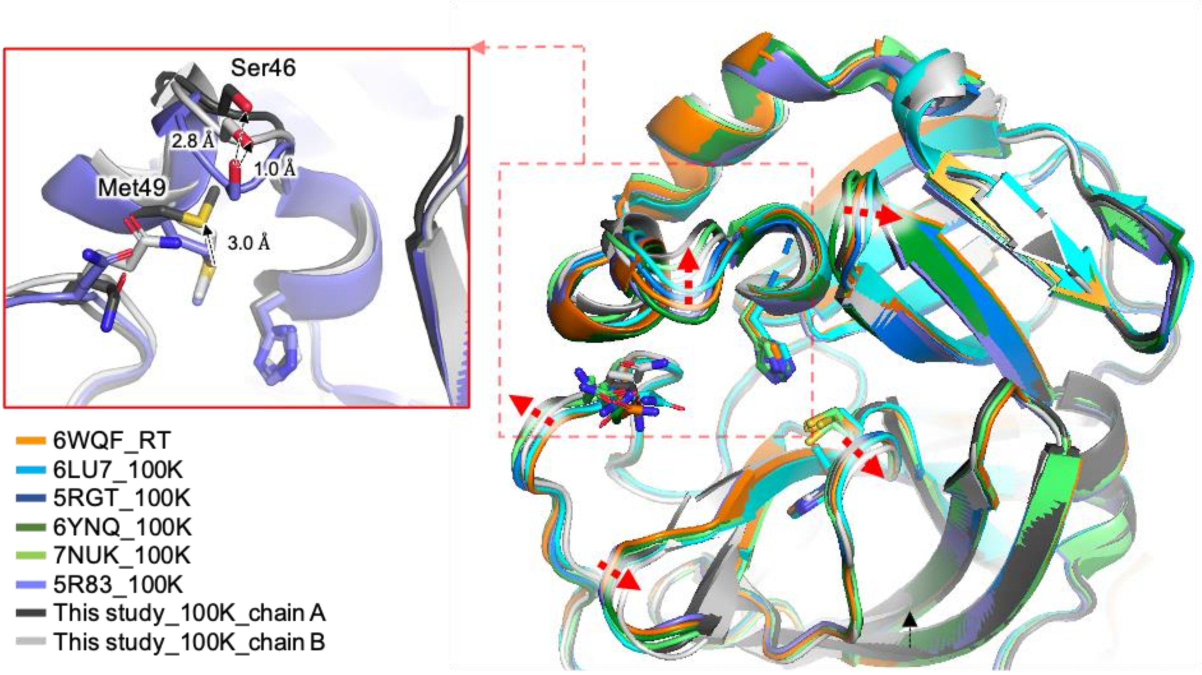


**Figure S1**

Comparison of the active site cavity conformations of different 3CL$^{pro}$ structures. The protein structures are shown in cartoon representation, with selected active site residues shown in a stick representation. The distance is indicated by dashed arrows and corresponding residues are highlighted.


**6WQF– Kneller et al 2020 (https://doi.org/10.1038/s41467-020-16954-7)**
**6LU7– Jin et al 2020 (https://doi.org/10.1038/s41586-020-2223-y)**
**5RGT– Zaidman et al 2021 (https://doi.org/10.1016/j.chembiol.2021.05.018)**
**6YNQ– Gunther et al., 2021 (https://doi.org/10.1126/science.abf7945)**
**7NUK– Sutanto et al., 2021 (https://doi.org/10.1002/anie.202105584)**
**5R83– Douangamath et al., 2020 (https://doi.org/10.1038/s41467-020-18709-w)**

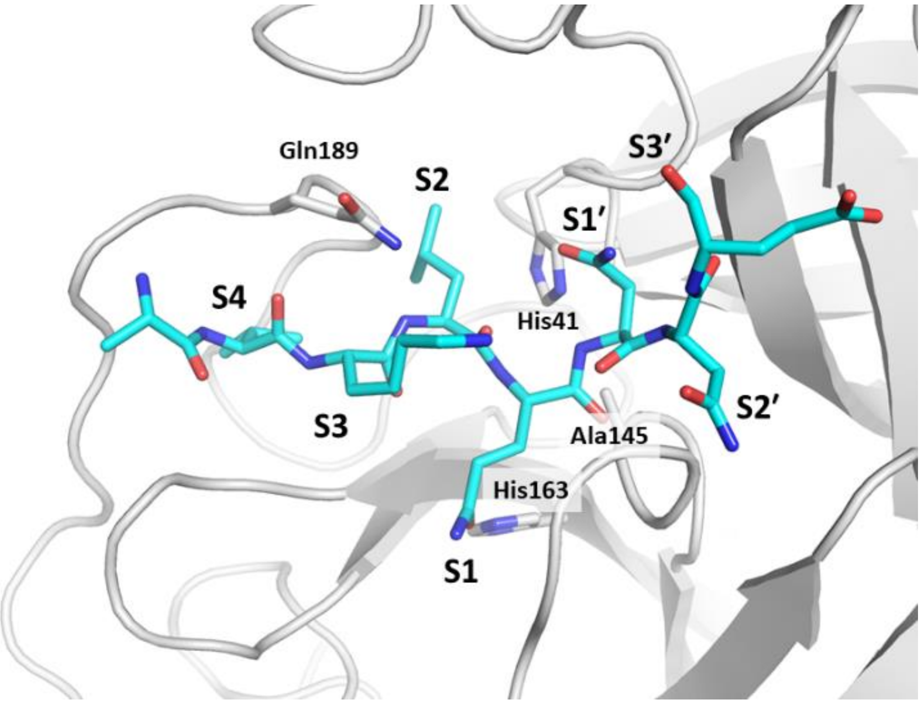


**Figure S2**

A substrate peptide bound to a catalytically inactive mutant of $3CL^{pro}$ (PDB entry 7mgr). The protein is shown in cartoon representation in grey, and the substrate peptide is shown in stick representations in blue. Stick models show C (cyan), N (blue) and O (red) atoms.

**Table S1. Data collection and refinement statistics.**

| | cpd-1 | cpd-2 | cpd-3 | cpd-4 | cpd-5 | cpd-6 | cpd-7 | cpd-8 | cpd-9 | cpd-10 | cpd-11 | cpd-12 | cpd-13 | cpd-14 | cpd-15 | cpd-16 | cpd-17 | cpd-18 | cpd-19 | cpd-20 | cpd-21 | cpd-22 | cpd-23 | cpd-24 | cpd-25 | cpd-26 | cpd-27 | cpd-28 | cpd-29 |
|---|---|---|---|---|---|---|---|---|---|---|---|---|---|---|---|---|---|---|---|---|---|---|---|---|---|---|---|---|---|
| **Data collection*** | | | | | | | | | | | | | | | | | | | | | | | | | | | | | |
| PDB code | 7GRE | 7GRF | 7GRG | 7GRH | 7GRI | 7GRJ | 7GRK | 7GRL | 7GRM | 7GRN | 7GRO | 7GRP | 7GRQ | 7GRR | 7GRS | 7GRT | 7GRU | 7GRV | 7GRW | 7GRX | 7GRY | 7GRZ | 7GR0 | 7GR1 | 7GR2 | 7GR3 | 7GR4 | 7GR5 | 7GR6 |
| Resolution range (Å) | 56.82-1.66 (1.69-1.66)** | 56.68-1.84 (1.88-1.84) | 56.81-1.54 (1.55-1.54) | 56.94-1.87 (1.91-1.87) | 57.09-1.79 (1.82-1.79) | 46.36-1.74 (1.77-1.74) | 56.25-1.80 (1.83-1.80) | 56.77-1.68 (1.7-1.68) | 56.65-1.70 (1.73-1.70) | 56.46-1.92 (1.97-1.92) | 56.68-1.55 (1.57-1.55) | 56.06-1.56 (1.58-1.56) | 56.70-1.67 (1.70-1.67) | 52.04-1.68 (1.71-1.68) | 45.93-1.47 (1.49-1.47) | 56.07-1.71 (1.75-1.71) | 56.71-1.81 (1.84-1.81) | 56.82-1.92 (1.96-1.92) | 56.94-1.92 (1.96-1.92) | 56.08-1.55 (1.57-1.55) | 56.64-1.54 (1.56-1.54) | 46.32-1.86 (1.89-1.86) | 55.81-1.69 (1.71-1.69) | 71.51-1.74 (1.78-1.74) | 56.77-1.77 (1.80-1.77) | 56.21-1.89 (1.94-1.89) | 56.85-1.86 (1.93-1.86) | 56.94-1.88 (1.95-1.88) | 56.16-1.62 (1.68-1.62) |
| Space group | $P2_12_12_1$ | $P2_12_12_1$ | $P2_12_12_1$ | $P2_12_12_1$ | $P2_12_12_1$ | $P2_12_12_1$ | $P2_12_12_1$ | $P2_12_12_1$ | $P2_12_12_1$ | $P2_12_12_1$ | $P2_12_12_1$ | $P2_12_12_1$ | $P2_12_12_1$ | $P2_12_12_1$ | $P2_12_12_1$ | $P2_12_12_1$ | $P2_12_12_1$ | $P2_12_12_1$ | $P2_12_12_1$ | $P2_12_12_1$ | $P2_12_12_1$ | $P2_12_12_1$ | $P2_12_12_1$ | $P2_12_12_1$ | $P2_12_12_1$ | $P2_12_12_1$ | $P2_12_12_1$ | $P2_12_12_1$ | $P2_12_12_1$ |
| Cell dimensions a, b, c (Å) | 67.79 99.93 104.20 | 67.71 99.47 103.64 | 67.78 100.18 104.17 | 67.97 101.02 104.27 | 68.1 101.23 104.7 | 68.03 100.96 104.38 | 67.83 100.65 104.01 | 67.78 99.57 103.9 | 67.61 99.63 103.81 | 67.31 99.63 103.69 | 67.62 99.49 103.93 | 67.75 99.82 103.89 | 67.73 99.35 103.68 | 67.77 100.55 104.08 | 67.64 99.77 103.48 | 67.78 99.79 103.54 | 67.68 99.44 103.91 | 67.76 100.04 104.28 | 67.94 100.67 104.39 | 67.87 99.57 103.83 | 67.68 99.16 103.50 | 67.85 100.65 104.34 | 67.53 99.15 103.22 | 67.60 99.03 103.36 | 67.75 99.52 104 | 67.25 99.17 102.39 | 67.74 99.96 104.54 | 67.92 99.96 104.43 | 67.99 99.62 104.44 |
| α=β=γ (°) | 90 | 90 | 90 | 90 | 90 | 90 | 90 | 90 | 90 | 90 | 90 | 90 | 90 | 90 | 90 | 90 | 90 | 90 | 90 | 90 | 90 | 90 | 90 | 90 | 90 | 90 | 90 | 90 | 90 |
| Total reflections | 850712 (39169) | 581538 (26315) | 1169474 (57088) | 575937 (29190) | 712847 (36987) | 672153 (32279) | 712405 (35462) | 936775 (40174) | 902494 (44166) | 536788 (30350) | 958735 (43164) | 929045 (45450) | 801741 (36103) | 799019 (37171) | 1128370 (54773) | 705856 (30303) | 780617 (40678) | 684097 (35913) | 5530002 (27680) | 1131064 (55998) | 1004192 (43267) | 837150 (134943) | 923113 (44141) | 669878 (28509) | 821338 (41410) | 440297 (20920) | 622297 (32253) | 556557 (28282) | 1066123 (51031) |
| Unique reflections | 62374 (3120) | 42769 (2139) | 85975 (4300) | 42625 (2131) | 51558 (2579) | 51113 (2557) | 52022 (2601) | 68538 (3428) | 66556 (3329) | 41138 (2057) | 70623 (3531) | 67649 (3382) | 60381 (3020) | 58288 (2914) | 83806 (4191) | 52132 (2608) | 56547 (2827) | 49742 (2487) | 41004 (2050) | 84285 (4214) | 73820 (3690) | 116173 (18522) | 67607 (3380) | 49438 (2472) | 59683 (2984) | 34351 (1719) | 46079 (2304) | 40569 (2029) | 77255 (3863) |
| Mean I / σ (I) | 9.9 (1.7) | 8.8 (1.7) | 14.9 (1.4) | 8.3 (1.7) | 11.1 (1.6) | 10.7 (1.5) | 10.6 (1.6) | 11.9 (1.6) | 14.2 (1.5) | 6.6 (1.5) | 11.6 (1.7) | 13.7 (1.8) | 7.4 (1.7) | 13.1 (1.8) | 15.6 (1.8) | 7.3 (1.7) | 8.8 (1.6) | 9.6 (1.5) | 7.4 (1.5) | 12.4 (1.4) | 12.7 (1.7) | 11.36 (1.08) | 10.9 (1.6) | 9.5 (1.6) | 9.8 (1.6) | 7.9 (1.6) | 10.7 (1.4) | 8.2 (1.5) | 16.8 (1.6) |
| Wilson B-factor | 35.08 | 38.05 | 31.77 | 46.42 | 39.24 | 43.216 | 36.57 | 34.04 | 24.17 | 43.107 | 33.62 | 33.51 | 32.15 | 37.95 | 28.82 | 38.668 | 35.63 | 41.27 | 41.64 | 30.34 | 31.23 | 40.97 | 34.70 | 34.16 | 32.65 | 35.20 | 41.74 | 42.968 | 31.97 |
| R-meas | 0.22 (2.06) | 0.29 (2.08) | 12.4 (2.13) | 0.21 (2.11) | 0.22 (2.15) | 0.18 (2.46) | 0.21 (2.18) | 0.18 (2.19) | 0.14 (2.07) | 0.38 (2.31) | 0.19 (2.18) | 0.14 (1.78) | 0.30 (2.21) | 0.15 (1.72) | 0.16 (1.70) | 0.27 (2.06) | 0.31 (2.27) | 0.2 (2.36) | 0.27 (2.32) | 0.17 (2.16) | 0.16 (1.83) | 0.12 (1.86) | 0.17 (2.32) | 0.38 (1.96) | 0.35 (2.27) | 0.38 (2.75) | 0.19 (2.16) | 0.22 (2.18) | 0.11 (1.88) |
| CC1/ 2 (%) | 99.7 (39.7) | 99.6 (52.0) | 99.9 (53.8) | 99.7 (42.5) | 99.7 (51.1) | 99.8 (64.4) | 99.8 (54.5) | 99.8 (48.0) | 99.9 (60.7) | 99.0 (33.5) | 99.8 (48.6) | 99.9 (57.0) | 99.5 (39.8) | 99.9 (66.7) | 99.9 (63.8) | 99.6 (51.8) | 99.6 (49.4) | 99.7 (46.9) | 99.4 (50.3) | 99.8 (55.2) | 99.9 (61.1) | 99.3 (70.2) | 99.8 (48.6) | 99.0 (60.9) | 99.5 (46.4) | 98.7 (98.9) | 99.8 (51.4) | 99.6 (47.8) | 99.9 (63.8) |
| Completeness spherical (%)*** | 74.6 (15.5) | 69.4 (12.4) | 81.0 (24.1) | 71.0 (13.5) | 74.9 (15.0) | 68.6 (12.2) | 78.1 (16.7) | 84.5 (26.3) | 96.2 (27.2) | 76.6 (18.3) | 68.9 (14.2) | 67.4 (11.3) | 74.0 (18.7) | 71.2 (13.0) | 70.0 (13.8) | 68.5 (12.1) | 88.2 (29.2) | 90.7 (38.3) | 73.8 (17.0) | 82.5 (24.2) | 70.8 (12.6) | 99.3 (98.1) | 86.1 (28.7) | 69.40 (14.0) | 86.7 (26.7) | 61.5 (13.8) | 76.8 (18.0) | 69.7 (13.7) | 85.7 (29.5) |
| Completeness ellipsoidal (%) | 91.3 (46.6) | 93.8 (62.9) | 94.4 (64.8) | 92.8 (56.9) | 93.8 (59.4) | 91.3 (50.9) | 94.4 (57.3) | 91.8 (43.4) | 86.2 (54.6) | 92.3 (68.2) | 91.9 (48.8) | 94.8 (64.3) | 90.4 (39.2) | 94.6 (64.0) | 95.3 (66.9) | 91.9 (54.5) | 93.8 (46.8) | 93.9 (50.6) | 93.6 (62.2) | 94.0 (58.5) | 95.2 (65.0) | - | 92.5 (48.3) | 93.40 (56.1) | 94.2 (48.0) | 79.8 (50.7) | 93.7 (57.4) | 93.3 (70.3) | 94.6 (63.5) |
| Multiplicity | 13.6 (12.6) | 13.6 (12.3) | 13.6 (13.3) | 13.5 (13.7) | 13.8 (14.3) | 13.2 (12.6) | 13.7 (13.6) | 13.7 (11.7) | 13.6 (13.3) | 13.0 (14.8) | 13.6 (12.2) | 13.7 (13.4) | 13.3 (12.0) | 13.7 (12.8) | 13.5 (13.1) | 13.5 (11.6) | 13.8 (14.4) | 13.8 (14.4) | 13.5 (13.5) | 13.4 (13.3) | 13.6 (11.7) | 7.2 (7.2) | 13.7 (13.1) | 13.5 (11.5) | 13.8 (13.9) | 12.8 (12.2) | 13.5 (14.0) | 13.7 (13.9) | 13.8 (13.2) |
| **Refinement** | | | | | | | | | | | | | | | | | | | | | | | | | | | | | |
| Reflections used in refinement | 62370 | 42759 | 85946 | 42624 | 51554 | 50966 | 52012 | 68505 | 66548 | 40980 | 70608 | 67641 | 60376 | 58285 | 83797 | 52109 | 56541 | 49735 | 40997 | 84280 | 73816 | 58522 | 67558 | 49437 | 59678 | 34289 | 46076 | 40565 | 77251 |
| $R_{work}$/ $R_{free}$ | 0.21/ 0.25 | 0.20/ 0.24 | 0.20/ 0.24 | 0.20/ 0.25 | 0.21/ 0.26 | 0.22/ 0.27 | 0.20/ 0.24 | 0.20/ 0.23 | 0.20/ 0.23 | 0.25/ 0.30 | 0.19/ 0.23 | 0.19/ 0.23 | 0.19/ 0.23 | 0.21/ 0.25 | 0.19/ 0.22 | 0.22/ 0.26 | 0.19/ 0.24 | 0.19/ 0.24 | 0.19/ 0.24 | 0.19/ 0.22 | 0.18/ 0.22 | 0.21/ 0.25 | 0.19/ 0.22 | 0.20/ 0.25 | 0.19/ 0.24 | 0.23/ 0.28 | 0.23/ 0.28 | 0.20/ 0.25 | 0.19/ 0.23 |
| Number of non-hydrogen atoms | 5324 | 5142 | 5313 | 5063 | 5110 | 5091 | 5176 | 5258 | 5118 | 5074 | 5394 | 5362 | 5241 | 4970 | 5574 | 5006 | 5242 | 5132 | 5101 | 5387 | 5336 | 5112 | 5208 | 5193 | 5276 | 5088 | 5122 | 4945 | 5275 |
| macromolecules | 4722 | 4687 | 4730 | 4694 | 4664 | 4691 | 4670 | 4721 | 4686 | 4658 | 4777 | 4724 | 4693 | 4706 | 4734 | 4682 | 4718 | 4697 | 4654 | 4739 | 4749 | 4663 | 4732 | 4691 | 4705 | 4709 | 4684 | 4676 | 4707 |
| ligands | 58 | 54 | 36 | 89 | 37 | 49 | 35 | 46 | 66 | 59 | 76 | 45 | 47 | 45 | 129 | 56 | 52 | 57 | 70 | 82 | 54 | 64 | 41 | 54 | 52 | 35 | 36 | 83 | 69 |
| solvent | 544 | 401 | 547 | 280 | 409 | 351 | 471 | 491 | 366 | 357 | 541 | 593 | 501 | 219 | 711 | 268 | 472 | 378 | 377 | 566 | 533 | 385 | 435 | 448 | 519 | 344 | 402 | 186 | 499 |
| RMS(bonds) (Å) | 0.008 | 0.004 | 0.007 | 0.006 | 0.004 | 0.004 | 0.008 | 0.039 | 0.020 | 0.009 | 0.009 | 0.022 | 0.010 | 0.011 | 0.009 | 0.019 | 0.097 | 0.012 | 0.005 | 0.008 | 0.015 | 0.005 | 0.011 | 0.066 | 0.010 | 0.003 | 0.004 | 0.010 | 0.013 |
| RMS(angles) (°) | 0.96 | 0.67 | 0.98 | 0.86 | 0.61 | 0.66 | 1.03 | 1.22 | 1.04 | 1.08 | 1.08 | 1.79 | 1.17 | 1.17 | 1.06 | 0.89 | 3.37 | 1.29 | 0.79 | 0.97 | 1.43 | 0.77 | 1.22 | 2.71 | 1.09 | 0.52 | 1.02 | 1.10 | 1.40 |
| Ramachandran favored (%) | 97.86 | 97.51 | 98.19 | 97.35 | 98.16 | 97.84 | 97.84 | 97.68 | 98.16 | 97.16 | 98.52 | 98.51 | 98.50 | 97.82 | 98.03 | 98.00 | 98.51 | 97.84 | 98.16 | 97.84 | 98.01 | 97.83 | 98.68 | 98.84 | 98.02 | 96.86 | 98.17 | 96.01 | 98.17 |
| Ramachandran allowed (%) | 2.14 | 2.49 | 1.81 | 2.65 | 1.84 | 2.16 | 2.16 | 2.32 | 1.84 | 2.84 | 1.48 | 1.49 | 1.50 | 2.18 | 1.97 | 2.00 | 1.49 | 2.16 | 1.84 | 2.16 | 1.99 | 2.17 | 1.32 | 1.16 | 1.98 | 3.14 | 1.83 | 3.99 | 1.83 |
| Ramachandran outliers (%) | 0.00 | 0.00 | 0.00 | 0.00 | 0.00 | 0.00 | 0.00 | 0.00 | 0.00 | 0.00 | 0.00 | 0.00 | 0.00 | 0.00 | 0.00 | 0.00 | 0.00 | 0.00 | 0.00 | 0.00 | 0.00 | 0.00 | 0.00 | 0.00 | 0.00 | 0.00 | 0.00 | 0.00 | 0.00 |
| Clashscore | 4.02 | 4.26 | 3.72 | 5.59 | 2.90 | 4.49 | 4.41 | 3.71 | 3.19 | 7.39 | 3.95 | 4.02 | 5.23 | 5.65 | 4.88 | 4.06 | 5.82 | 6.49 | 4.93 | 2.83 | 3.16 | 3.33 | 4.13 | 3.30 | 5.00 | 4.80 | 6.11 | 6.71 | 4.78 |
| Average B-factor | 22.00 | 28.20 | 25.40 | 35.80 | 29.04 | 34.09 | 27.97 | 24.44 | 28.44 | 30.21 | 25.70 | 24.48 | 20.81 | 28.15 | 23.67 | 27.15 | 24.89 | 32.21 | 36.26 | 25.97 | 24.44 | 38.24 | 28.17 | 25.52 | 21.72 | 19.49 | 33.75 | 38.80 | 28.18 |
| macromolecules | 21.12 | 27.72 | 24.52 | 35.25 | 28.55 | 33.95 | 27.31 | 23.55 | 27.55 | 30.28 | 24.56 | 23.32 | 19.95 | 27.94 | 21.81 | 26.63 | 24.02 | 31.70 | 35.64 | 24.62 | 24.17 | 37.70 | 27.33 | 24.80 | 20.78 | 19.42 | 33.42 | 38.50 | 27.02 |
| ligands | 33.19 | 41.67 | 34.50 | 62.79 | 39.31 | 42.38 | 35.02 | 35.65 | 57.47 | 34.54 | 46.07 | 40.07 | 31.04 | 43.57 | 44.11 | 49.37 | 40.19 | 43.58 | 51.36 | 40.14 | 45.30 | 46.57 | 33.39 | 38.21 | 32.81 | 27.24 | 44.27 | 59.70 | 49.64 |
| solvent | 28.41 | 32.01 | 32.43 | 36.40 | 33.70 | 34.73 | 33.97 | 31.92 | 34.58 | 28.50 | 32.92 | 32.52 | 27.92 | 29.38 | 32.30 | 31.45 | 31.84 | 36.80 | 41.05 | 35.22 | 34.79 | 43.30 | 36.82 | 31.52 | 29.19 | 19.72 | 36.66 | 36.93 | 36.18 |

*Data processing statistics are reported with Friedel pairs separated.

**Statistics for the highest-resolution shell are shown in parentheses.

***With the exception of the cpd-22, which was processed by XDS, datasets were processed and elliptically truncated using the autoPROC STARANISO option.

**Table S2**
SMILES codes of $3CL^{pro}$ fragment hits.

| Compound | SMILES code | Compound | SMILES code |
|---|---|---|---|
| 1 | FC(c1cc(-c2ccncc2)n[nH]1)(F)F | 16 | O=C(c1ccccc1)NCc(cc1)cc2c1OCC2 |
| 2 | Nc1cc(Br)cnc1 | 17 | C[n](c(N)c1)nc1-c(cc1)ccc1Cl |
| 3 | Nc(c(Cl)cnc1)c1Cl | 18 | O=C1NC(c(cccc2)c2F)SC1 |
| 4 | Oc1cc(Cl)cnc1 | 19 | CC(C(Nc1cc(Cl)cc(Cl)c1)=O)O |
| 5 | CC(c1ccn[nH]1)=O.Cl | 20 | O=C(CCC1=O)N1c(ccc(F)c1)c1F |
| 6 | OCc1c[s]c(cc2)c1cc2Cl | 21 | O=C(CCC1=O)N1c1cc(Cl)cc(Cl)c1 |
| 7 | OCc1cc(F)cc2c1OCOC2 | 22 | CN(C)C(COc1cc2ccccc2cc1)=O |
| 8 | CC(CC[n](nc1Br)nc1Br)=O | 23 | O=C(c1ncccc1)c1nc2ccccc2cc1 |
| 9 | CC(Nc(cc1)cc(CCC2)c1C2=O)=O | 24 | N#CCC(NC1CCCCC1)=O |
| 10 | FC(c(cc1)cc2c1[nH]c([C@@H]1CNCC1)n2)(F)F | 25 | OC(c1cc(-c2cnccc2)ccc1)=O |
| 11 | CC(Nc(cc1)c(C(F)(F)F)cc1C#N)=O | 26 | NCc(cc1)cnc1-c1ccccc1 |
| 12 | NCc(cc1)cc2c1OCC2 | 27 | CC(N12)=CSC1=NC(C=O)=CC2=O |
| 13 | Cc1c(CNC(c2ccccc2)=O)[s]cc1 | 28 | O=C(c(cc(cc1)F)c1N1)C1=O |
| 14 | OC(c1cncc(C#CCC2CCCCC2)c1)=O | 29 | O=C(c(c(c(N1)c(cc2)Cl)c2Cl)C1=O |
| 15 | CNCc(cc1)ccc1-c1ccc[s]1 | | |
